# Soft Sensing of Intracellular States for CHO Cell Bioprocessing with Ensemble Kalman Filters

**DOI:** 10.64898/2026.05.28.728559

**Authors:** Luxi Yu, Antonio del Rio Chanona, Cleo Kontoravdi

## Abstract

In biotherapeutic manufacturing, product quality such as glycosylation profile is typically assessed only after harvest, limiting opportunities for corrective action during cell culture operation. Intracellular nucleotide sugar donors (NSD) directly determine glycosylation outcomes but are rarely measured, even offline, due to analytical complexity and process disruption. As a result, quality-related decisions remain constrained to fixed operating strategies. This work introduces a model-based soft sensing framework to infer NSD concentrations from readily available extracellular measurements. A Bayesian state estimation approach based on the Ensemble Kalman Filter (EnKF) is developed to reconstruct unmeasured intracellular states during CHO cell culture. An imperfect kinetic process model is combined with noisy extracellular measurements, explicitly accounting for process variability and measurement uncertainty through ensemble-based propagation and updates. The framework is validated using four independent experiments with distinct feeding perturbations that are not used for model calibration. Although the open-loop model exhibited substantial mismatch for both extracellular metabolites and intracellular NSDs, EnKF assimilation of extracellular measurements corrected key metabolic profiles. Building on these corrected extracellular dynamics, the EnKF demonstrated robust estimation of a growth-determining amino acid, asparagine, from correlated extracellular states. Based on the improved extracellular and amino acid estimates, the framework further enabled reliable inference of intracellular NSDs across all experiments.

## 1 Introduction

Accurate and timely information on key process variables is essential for effective monitoring and control of biotherapeutic manufacturing processes. In current industrial practice, routine online monitoring is largely confined to physical operating conditions such as temperature, pH, and dissolved oxygen tension, while most biologically relevant information is obtained through offline analysis. Extracellular process variables, including nutrient and metabolite concentrations, and product titre, are typically measured with some delay, although advances in process analytical technologies such as Raman spectroscopy are expected to increase the availability of (near) real-time extracellular measurements [1–8]. In contrast, intracellular metabolites are rarely measured in manufacturing environments, owing to experimental complexity, analytical cost, and disruption to the process [9, 10]. However, certain intracellular metabolites are directly correlated with key process outputs. Specifically, nucleotide sugar donors (NSDs) serve as essential substrates for *N*-linked glycosylation, a critical quality attribute that strongly influences the safety, efficacy, and pharmacokinetics of recombinant protein products [11, 12]. For example, the presence of galactose residues influences complement-dependent cytotoxicity responses [13], terminal sialylation enhances anti-inflammatory properties [14] and prolongs serum half-life [15]. In contrast, fucosylation reduces antibody-dependent cellular cytotoxicity activity [16], high-mannose type and glycans with terminal *N*-acetylglucosamine residues are found to exhibit more rapid clearance than more fully glycosylated antibodies [17–19]. Glycosylation profiles are typically assessed only at harvest, with analytical results available after the completion of the cell culture run. As a consequence, product quality is evaluated retrospectively without feedback during operation, and no corrective intervention can be made once deviations from the desired glycan profile are identified. Moreover, in the absence of information on the dynamic intracellular availability of NSDs, it is not possible to anticipate quality-relevant shifts in glycosylation during the run, even when extracellular measurements indicate metabolic changes. This lack of transparency therefore limits quality-oriented decision-making to fixed operating strategies and *post hoc* analysis.

Mechanistic modelling has played a central role in understanding NSD contribution to glycosylation and in establishing NSDs as a key link between extracellular process conditions and product quality. Early kinetic frameworks explicitly represented NSD synthesis, transport, and consumption within the Golgi apparatus. For example, del Val *et al*. developed a mechanistic description of intracellular NSD availability relevant to glycosylation [20], while Jedrzejewski *et al*. proposed an integrated model linking extracellular metabolite uptake rates to intracellular NSD production and, in turn, glycan distribution on the recombinant protein product [21]. Building on these foundational work, subsequent efforts focused on developing predictive models for understanding how variations in the culture environment influence NSD availability and glycosylation outcomes. Key studies investigated the effects of operating conditions such as pH [22, 23], dissolved oxygen tension [23], osmolarity [23], temperature shifts [24], and supplementation strategies such as NSD precursor feeding [22, 25], capturing culture behaviour in kinetic modelling glycosylation frameworks. While these kinetic models provide valuable mechanistic insight, extrapolation of these models has been shown to be poorly predictive of cellular regulation events [26], owing to their reliance on fixed parameter values and predefined kinetic structures. Consequently, their applicability across different cell lines, operating conditions, and products is limited, and their robustness is reduced in the presence of process uncertainty and batch-to-batch variability.

In an effort to improve flexibility, alternative modelling strategies have sought to reduce reliance on detailed intracellular kinetics by exploiting stoichiometric constraints. Two-step frameworks have been proposed in which extracellular metabolite measurements are used to estimate intracellular NSD-related fluxes via constraint-based flux balance analysis, followed by coupling to either kinetic [27] or artificial neural network models [28] for glycosylation prediction. Although this approach enables indirect inference of NSDs from extracellular data and alleviates some parametrisation challenges, it relies on pseudo-steady-state assumptions and does not explicitly capture the transient dynamics of intracellular NSD pools over the course of a fed-batch culture.

Across existing modelling approaches, a persistent limitation is the lack of systematic treatment of uncertainty and variability inherent to bioprocesses, including unmodelled dynamics, measurement noise, and batch-to-batch heterogeneity. These challenges become particularly problematic when the quantities of interest are intracellular and difficult to measure, as is the case for NSD concentrations. In such settings, process models alone are insufficient to recover dynamic state information, and systematic integration of model predictions with available measurements becomes necessary.

Model-based soft-sensing techniques address this need by integrating imperfect process models with sparse and noisy measurements within a state estimation framework, explicitly accounting for uncertainties. Classical nonlinear Kalman filtering methods, such as the Extended Kalman Filter (EKF), achieve this by linearising the process model around the current estimate and propagating error covariances accordingly [29]. Although fast and effective for mildly nonlinear systems, EKF performance degrades in the presence of strong nonlinearities and discontinuities [30], which are common characteristics of mammalian cell culture processes. The Unscented Kalman Filter (UKF) addresses this limitation of the EKF by approximating the propagation of state uncertainty through nonlinear dynamics using a set of deterministically chosen sigma points [31]. While avoiding explicit linearisation, the accuracy of the UKF depends on the selection of tuning parameters that govern the spread and weighting of these sigma points, often requiring careful calibration to ensure numerical stability and reliable estimation performance [32]. Alternatively, Particle Filters (PF) provide a non-parametric representation of uncertainty but are typically computationally demanding and prone to sample degeneracy in high-dimensional systems [33]. Moving horizon estimation offers a flexible optimisation-based formulation [29, 34, 35] but requires repeated solution of constrained nonlinear optimisation problems and explicit handling of arrival costs [36–38], limiting its practicality for real-time soft sensing in complex bioprocess models.

Here, we introduce the Ensemble Kalman Filter (EnKF) for soft-sensing of unmeasured intracellular states in mammalian cell culture processes. The EnKF was originally developed for large-scale state estimation problems in geosciences, where systems are highly nonlinear, partially observed, and subject to substantial uncertainty [39]. Its ensemble-based formulation represents state uncertainty through a collection of model realisations, which are propagated through the nonlinear process model to recover error statistics without explicit linearisation or storage of full covariance matrices [40]. This distinguishes the EnKF from classical nonlinear Kalman filtering approaches, which rely on local linearisation or deterministic approximations, as well as from particle filtering and optimisation-based methods, which can become computationally prohibitive in higher-dimensional settings. These properties make the EnKF particularly well suited for soft sensing in bioprocessing applications, where nonlinear dynamics, measurement sparsity, and process variability are inherent.

We have previously demonstrated the feasibility of estimating intracellular NSD concentrations *in silico* using an EnKF framework supported by comprehensive extracellular measurements, including amino acids [41]. In the present study, this framework is extended to a more realistic soft-sensing scenario and validated experimentally, under the assumption that only a limited set of routinely measured extracellular metabolites is accessible. Within this setting, both asparagine and intracellular NSDs are treated as unmeasured during filtering and are inferred solely through their coupling to observable extracellular states.

Independent fed-batch experiments employing distinct galactose and uridine feeding strategies are used for validation. Both asparagine and intracellular NSD concentrations are inferred dynamically using the EnKF and validated against independently measured offline data that are not assimilated by the filter. The results presented in this study show that the EnKF enables dynamic inference of unmeasured intracellular states from limited extracellular data, providing information that supports earlier, quality-relevant intervention during bioprocess operation.

## 2 Materials and Methods

### 2.1 Mechanistic Model and Experimental Data

#### 2.1.1 Mechanistic Model

The mechanistic model employed in this study comprises two coupled components representing extracellular cell culture behaviour and intracellular NSD metabolism, based on previously published work [25]. The cell culture submodel describes viable cell growth and death, extracellular metabolite uptake and secretion, and monoclonal antibody synthesis. The NSD module captures the synthesis of intracellular nucleotide sugar donors as a function of the specific growth rate, together with their transport into the Golgi apparatus, where they act as substrates for glycosylation reactions. The full set of model equations, state definitions, kinetic expressions, and nominal parameter values are provided in Supplementary Information A.

#### 2.1.2 Calibration Data

Model parameters were identified using five fed-batch experiments reported by Kotidis *et al*. (2019), generated from an IgG-producing CHO-T127 cell line operated under different galactose and uridine supplementation strategies [25]. All experiments were conducted in suspension culture using CD CHO medium at 36.5 °C and 5 % CO_2_, with orbital agitation at 150 rpm. Cultures were initiated at a seeding density of 2 × 10^5^ cells mL^-1^ in 500 mL vented Erlenmeyer flasks with a working volume of 100 mL and supplemented with 10 % (v/v) CD EfficientFeed C™ AGT™ on alternating days starting from Day 2. Manganese(II) chloride was added at seeding at a concentration of 1 *µM* in all fed-batch conditions to support *β*-1,4-galactosyltransferase activity. Five datasets were used exclusively for parametrisation of the mechanistic model prior to its integration with the EnKF. Detailed descriptions of the feeding strategies, sample preparation protocols, and analytical quantification procedures are provided in Supplementary Information Section B.

#### 2.1.3 Validation Experiment

The performance of the EnKF-based estimation framework was evaluated using four additional fed-batch experiments that were not used during model parametrisation. These experiments were conducted independently but employed the same CHO-T127 cell line, culture medium, and operating conditions as those used for model calibration, allowing validation to focus on the framework’s ability to generalise across feeding strategies rather than changes in process configuration. Each experiment applied a distinct galactose and uridine supplementation profile, selected to span a design space constrained by requirements on product titre and galactosylation quality, as reported in [26]. The corresponding feeding conditions are summarised in Table 1. The same analytical methods described in Supplementary Information B were used for all validation experiments.

**Table 1:** Galactose and uridine feeding concentrations (mM) applied in each experiment. P4 used for EnKF covariance tuning; P1-P3 used for validation.

| Exp. | Galactose (mM) |  |  |  | Uridine (mM) |  |  |  |
| --- | --- | --- | --- | --- | --- | --- | --- | --- |
|  | Day 4 | Day 6 | Day 8 | Day 10 | Day 4 | Day 6 | Day 8 | Day 10 |
| P1 | 79.4 | 15.4 | 11.0 | 248.3 | 15.9 | 3.1 | 2.2 | 49.7 |
| P2 | 4.3 | 168.3 | 37.7 | 11.4 | 0.9 | 33.7 | 7.5 | 2.3 |
| P3 | 5.2 | 3.1 | 235.3 | 249.9 | 1.0 | 0.6 | 47.1 | 50.0 |
| P4 | 21.9 | 6.4 | 233.5 | 4.0 | 4.4 | 1.3 | 46.7 | 0.8 |

### 2.2 Empirical observability analysis

Assessing the observability of unmeasured or partially measured states is essential for reliable state estimation in complex bioprocess systems. Classical observability analysis, developed for linear time-invariant systems and based on rank conditions of a state-space representation, has limited applicability to inherently nonlinear and high-dimensional biological models. Linearisation-based tests provide only local insight around a chosen operating point and fail to capture the global input–output sensitivity of nonlinear dynamics. Moreover, such rank-based criteria yield binary conclusions and do not reflect varying degrees of state inferability, while the analytical computation of Jacobians becomes impractical for complex bioprocess models.

To address these challenges, a simulation-based observability assessment based on the empirical observability Gramian was adopted. This approach avoids explicit linearisation and instead quantifies the sensitivity of measured outputs to perturbations in individual state variables [42]. Specifically, the nominal initial state vector *x*_0_ ∈ ℝ^*n*^ was first normalised to obtain dimensionless states, thereby mitigating numerical issues arising from large differences in state magnitudes. Each state component was then perturbed individually by a small symmetric offset *±ε*, yielding perturbed initial conditions 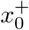 and 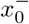. These states are converted back to their dimensional form and propagated forward in time using the mechanistic model, generating corresponding output trajectories *y*^+^(*t*) and *y*^−^(*t*). The outputs are subsequently normalised to obtain dimensionless trajectories 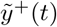 and 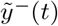. The influence of each state on the measured outputs was quantified through the dimensionless output difference

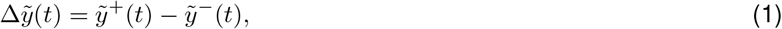

which was integrated over the simulation horizon *T*. The resulting diagonal entry of the empirical observability Gramian was computed as

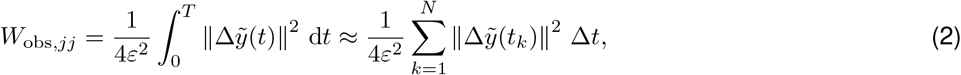

where Δ*t* denotes the fixed simulation time step and *N* Δ*t* = *T*. Larger values of *W*_obs,*jj*_ indicate that perturbations in the corresponding state lead to pronounced changes in the measured outputs, implying that the state is well observed under the available measurement configuration, whereas small or near-zero values indicate weak or negligible influence on the outputs.

### 2.3 Ensemble Kalman Filter

The EnKF is a Monte Carlo-based extension of the Kalman Filter designed for nonlinear and high-dimensional systems [43]. Instead of propagating state covariances analytically, the EnKF represents uncertainty using an ensemble of state trajectories that are evolved forward through the nonlinear process model and updated sequentially as measurements become available. The posterior state distribution is approximated through repeated prediction and measurement update steps, using a linear-Gaussian correction applied consistently across all ensemble members [44]. Although the Gaussian assumption in the update step may limit its accuracy for strongly non-Gaussian systems, the EnKF remains computationally efficient and is widely used in large-scale nonlinear estimation problems [40].

Figure 2 summarises the EnKF workflow implemented in this study. An ensemble of state vectors is initialised by sampling from a multivariate normal distribution,

**Figure 1:**
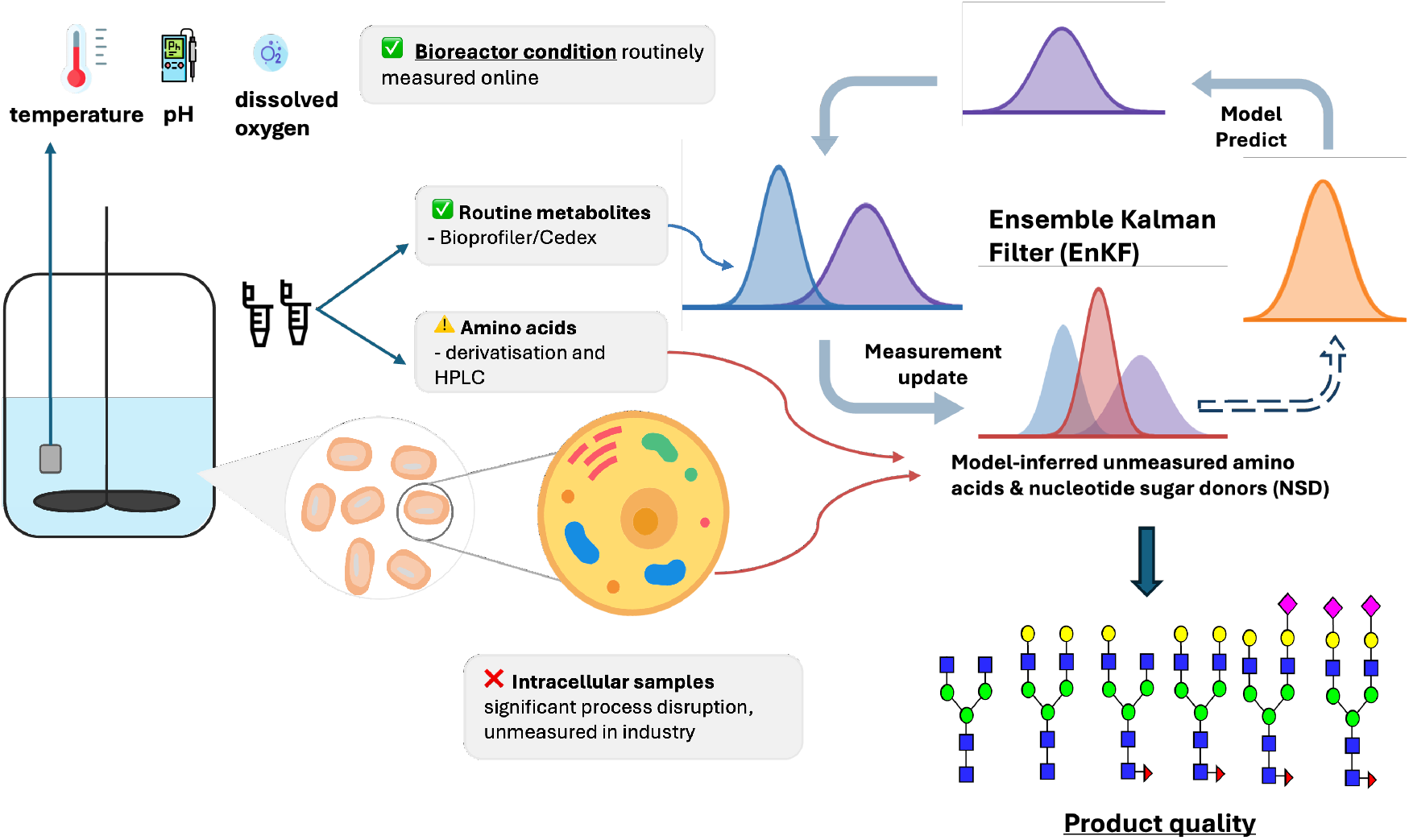
Schematic of EnKF-based state estimation in bioreactor operation, integrating routine extracellular measurements with a mechanistic model to infer unmeasured amino acids and intracellular states, which can be used for predicting product quality.

**Figure 2:**
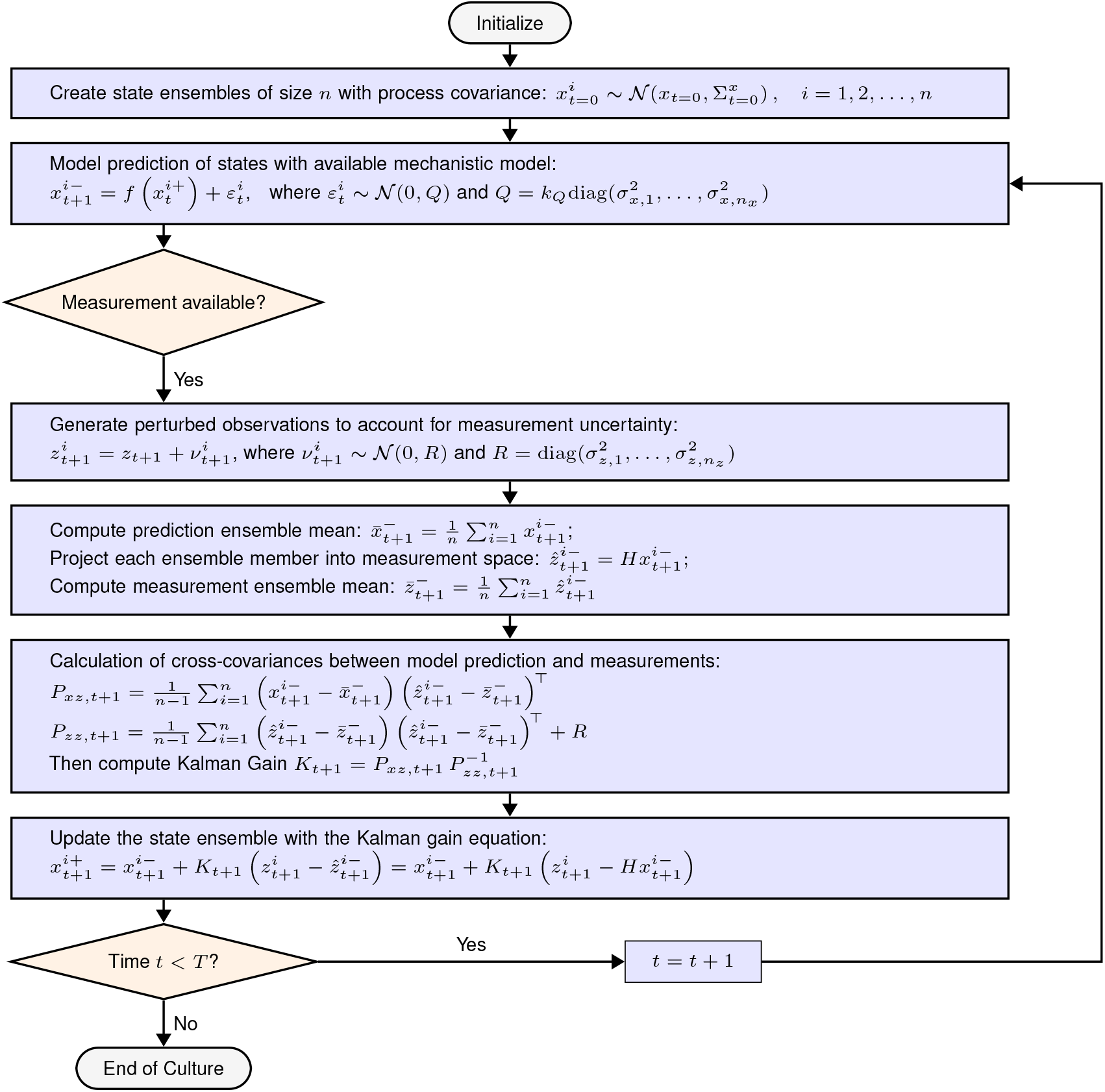
Schematic representation of the EnKF used for state estimation. State ensembles are propagated using the mechanistic model and updated when measurements are available. Measurement uncertainty is accounted for through ensemble perturbation, and state corrections are obtained from ensemble-derived covariances between model predictions and measurements.

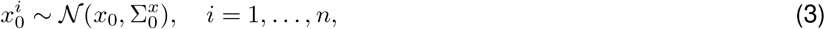

where *x*_0_ denotes the nominal initial condition and 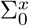 the associated initial ensemble covariance. An ensemble size of *n* = 100 was used, which was found to be sufficient for the 17-state system considered. Each scenario was simulated 10 times and the mean trajectory was reported, ensuring repeatability while preserving the stochastic nature of the EnKF. Owing to the slow dynamics of the bioprocess, this level of repeated simulation remains computationally affordable, even for real-time state estimation and update.

During the prediction step, each ensemble member is propagated forward using the nonlinear mechanistic model,

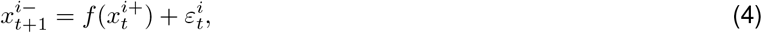

where *f* (*·*) denotes the system dynamics and the superscripts (+) and (−) indicate states after and before the measurement update, respectively. The additive process noise term is introduced to represent model mismatch and unmodelled dynamics, and is modelled as

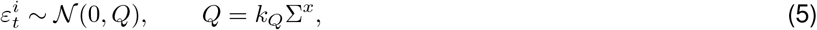

where 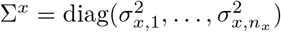 defines a fixed, state-wise variance structure and the scalar factor *k*_*Q*_ controls the overall magnitude of process noise while preserving relative uncertainty across states. The ensemble-based state covariance is estimated directly from ensemble statistics and evolves dynamically at each prediction and update step, whereas the additive process noise covariance *Q* is prescribed and held constant. This formulation was found to be effective in maintaining sufficient ensemble spread, particularly in systems with correlated states and partial observability [39].

In this study, the measurements *z*_*t*+1_ are treated as given experimental observations rather than as outputs of a generative measurement model. The relationship between system states and the measured variables is introduced through a linear observation operator,

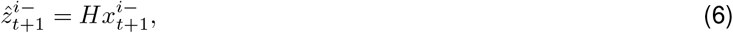

which maps the predicted state of ensemble member *i* into the measurement space. Here, 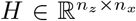 selects the measured extracellular variables and sets unmeasured components to zero. In practice, *H* is constructed as *H* = [*I* 0], such that measured states are directly observed, while unmeasured intracellular states are not directly corrected but may still be influenced indirectly through correlations captured by the ensemble.

Measurement uncertainty is characterised by the covariance matrix

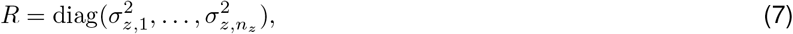

defined only for the measured variables. To incorporate this uncertainty within the EnKF framework, a stochastic update is employed by generating perturbed observations for each ensemble member,

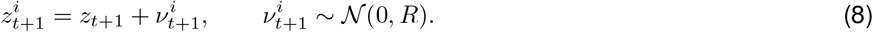

Following the prediction step, ensemble statistics are computed. The ensemble mean of the predicted states is given by

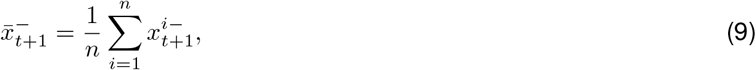

and the corresponding ensemble mean in measurement space is

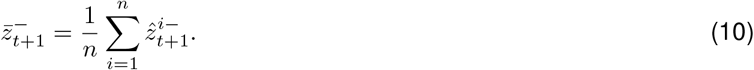

The coupling between state uncertainty and measurements is quantified using ensemble-based covariance estimates, computed directly from ensemble statistics as:

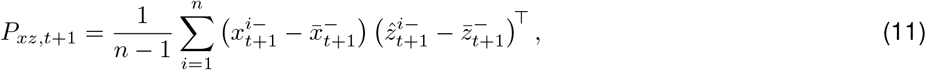

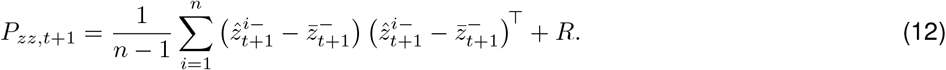

The Kalman gain is subsequently computed as

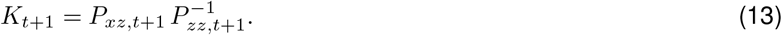

Each ensemble member is then updated according to

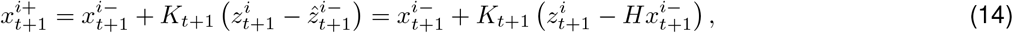

where the correction term represents the confidence between the experimental measurement and the model prediction. Although only a subset of states is directly measured, unmeasured states are adjusted implicitly through ensemble cross-covariances with the measured variables. The prediction and update steps are repeated sequentially until the end of culture. To ensure numerical stability and physical feasibility, ensemble members are constrained to remain non-negative following both prediction and update steps. The calibrated process and measurement noise covariances used in this study are reported in Supplementary Information C.

## 3 Results and Discussion

### 3.1 Empirical Observability Analysis

#### 3.1.1 Extracellular Metabolite Observability

Key extracellular metabolites including glucose, lactate, ammonia, and glutamine can be easily measured using standard analytical platforms such as the BioProfile (Nova Biomedical) or CedexBio Analyzer (Roche CustomBiotech), which are widely adopted in bioprocessing industries. Viable cell density and monoclonal antibody titre are also routinely measured. In contrast, asparagine cannot be monitored in real time and typically requires labour-intensive offline analysis based on amino acid derivatisation and chromatographic separation.

Empirical observability analysis shown in Figure 3 (a) reveals differences in how individual extracellular states influence measurable system outputs. Viable cell density, glucose, and ammonia exhibit consistently high Gramian scores, reflecting their strong dynamic coupling to the system through growth and death kinetics. Perturbations in these states propagate rapidly and produce measurable effects, resulting in high empirical observability. In contrast, antibody titre, lactate, and glutamine show lower observability scores, consistent with weaker dynamic coupling or limited sensitivity of the measured outputs to perturbations in these states. Among supplemented precursors for glycosylation, uridine displays substantially higher observability than galactose. This difference arises from model structure, where uridine uptake directly affects the specific death rate leading to rapid propagation of perturbations through the system, while galactose influences the dynamics more indirectly via modulation of glucose uptake, resulting in reduced output sensitivity.

**Figure 3:**
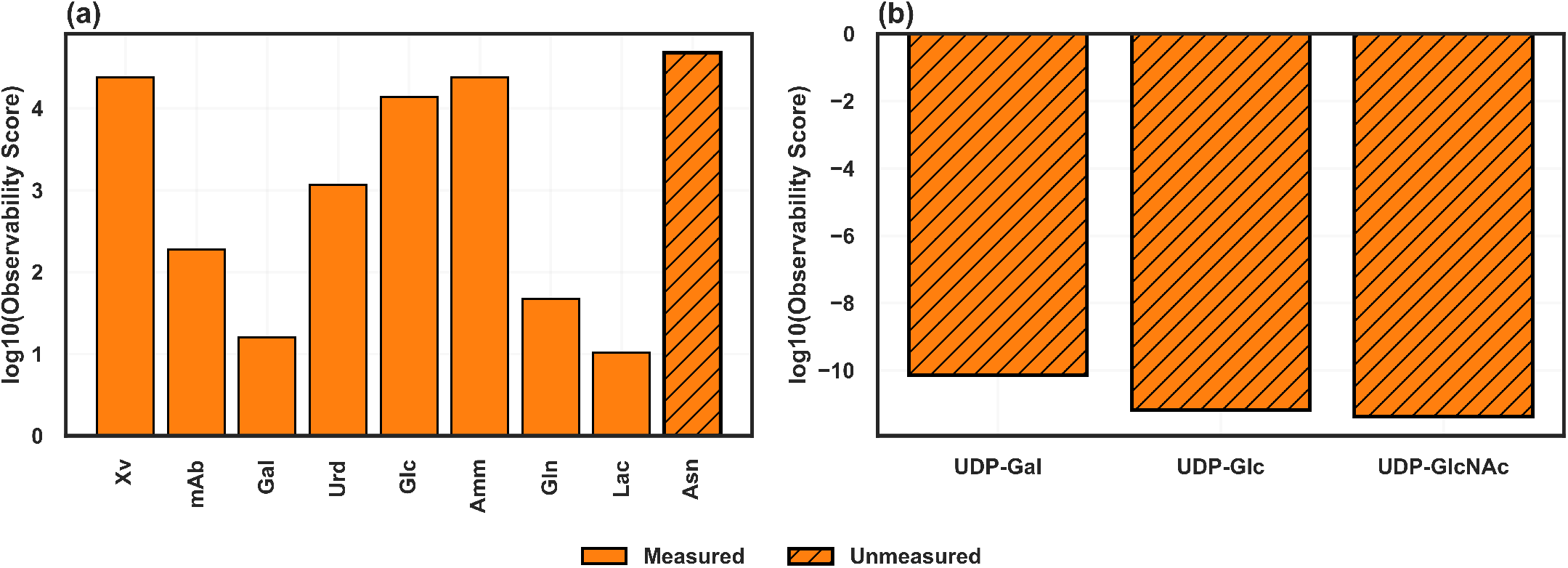
(a) Empirical observability scores for model state variables under the available measurement configuration. (b) Empirical observability scores for selected intracellular NSD (UDP-Gal, UDP-Glc, and UDP-GlcNAc) evaluated using the nonlinear observability Gramian.

Notably, asparagine exhibits one of the highest observability scores among all states, despite being unmeasured. In the model, asparagine is tightly coupled to the specific growth rate, which, in turn, governs viable cell density, protein synthesis, and the uptake and secretion of extracellular metabolites. Consequently, small perturbations in asparagine propagate and generate measurable effects on system outputs, leading to strong empirical observability.

#### 3.1.2 Intracellular NSD Observability

NSDs are synthesised intracellularly from extracellular metabolites such as glucose, glutamine, galactose, and uridine through a sequence of cytosolic enzymatic reactions, and are subsequently consumed in the Golgi apparatus during glycosylation. This system structure reflects an inherent biological property characterised by a strictly unidirectional information structure, in which extracellular states drive NSD synthesis, while variations in NSD concentrations do not propagate measurable effects back to the extracellular environment.

This structural property is reflected in the empirical observability analysis in Figure 3 (b), which confirms that NSDs are effectively unobservable under the given measurement configuration. Because the observability Gramian is computed using full nonlinear simulations, near-zero values indicate structural nonlinear unobservability rather than weak local sensitivity that could be recovered through higher-order effects. Under these conditions, no estimator can recover NSD states uniquely from measurements alone, regardless of filter type. Any estimation of NSD dynamics must therefore rely on model-based inference conditioned on upstream observable states, rather than direct correction from measurement residuals.

For structurally unobservable states, classical linearised estimators such as the EKF are not suitable. The EKF relies on Jacobian-based linearisation of both the process and measurement models to propagate state covariances and compute the correction Kalman gains [45]. When the unmeasured states do not influence the output measurements, the corresponding sensitivities vanish, leading to degenerate or ill-conditioned covariance updates. In practice, uncertainty in NSD states accumulates without meaningful correction, while linearisation errors introduced at each update step are not dissipated by measurement feedback, resulting in numerical instability or divergence.

By contrast, the EnKF provides a more suitable estimation framework for this class of problems. Instead of relying on Jacobian-based local linearisation and analytical covariance propagation as in EKF, the EnKF represents state uncertainty through an ensemble of full nonlinear system trajectories, with the sample covariance of the ensemble providing a Monte Carlo approximation of the evolving state uncertainty. This ensemble-based formulation ensures that uncertainty propagation remains consistent with the nonlinear process dynamics, allowing variability in observable extracellular metabolite states to be transmitted to intracellular NSD concentrations through mechanistic coupling, without imposing spurious linear sensitivity. While the EnKF does not overcome the structural unobservability of NSDs and the inferred posterior distributions remain dependent on model structure and uncertainty specification, it provides a numerically more robust method for constraining NSD evolution on available extracellular metabolite measurements.

### 3.2 EnKF Estimation of Measurable Extracellular States

Figure 4 compares EnKF-based state estimates of measurable extracellular variables with open-loop mechanistic model predictions, demonstrating consistently improved agreement with experimental data across all measured states. Discontinuities in the model trajectories arise from the bolus feeding strategy applied every two days, whereas the piecewise adjustments in the EnKF estimates reflect daily measurement updates.

**Figure 4:**
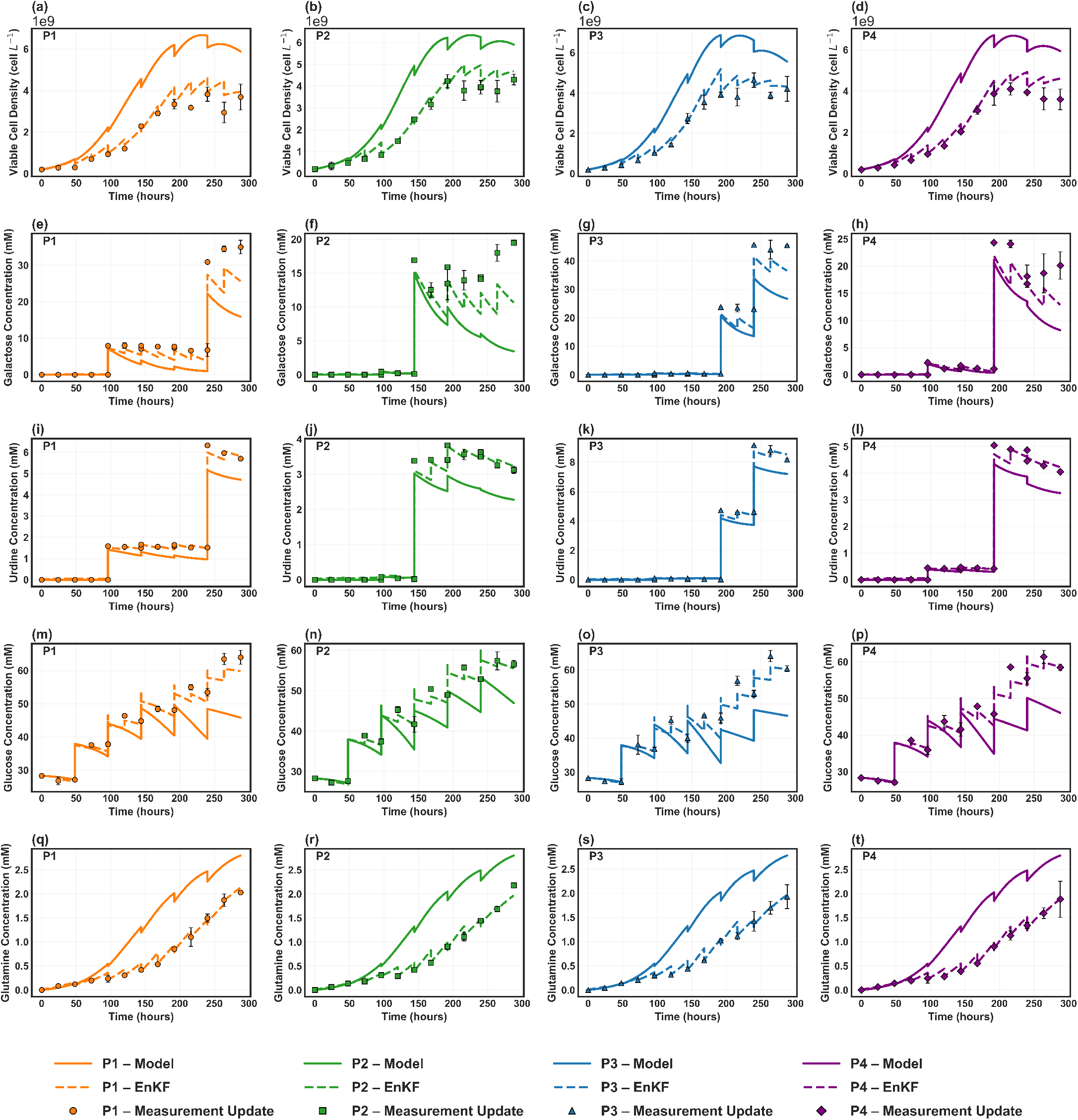
EnKF-based state estimation of measurable extracellular variables across four feeding strategies (columns). Only routinely available measurements are assimilated in the EnKF updates, including viable cell density, monoclonal antibody titre, galactose, uridine, glucose, glutamine, lactate, and ammonia. All other available data are reserved exclusively for validation and are not used during state estimation.

As shown in Figure 4 (a)-(d), the mechanistic model systematically overestimated viable cell density for all feeding strategies, even though it was parametrised using data from the same cell line. The consistent reduction in growth observed across all conditions, together with biological replicates and reasonable measurement uncertainty, suggests that this discrepancy is unlikely to be driven by noise. Instead, it more plausibly reflects genuine deviations from nominal model behaviour arising from biological variability or subtle differences in cultivation conditions. The EnKF effectively compensates for this mismatch by assimilating viable cell density measurements, recovering growth trajectories that closely follow the experimental profiles.

Galactose and uridine, which serve as key inputs to intracellular precursor synthesis pathways relevant to glycosylation, required careful tuning of process and measurement noise. As shown in Figure 4 (e)-(h), the EnKF estimates for galactose aligned closely with experimental measurements while avoiding overfitting of spurious fluctuations, particularly in regions where apparent concentration increases occurred in the absence of feeding events. This behaviour is consistent with known limitations of fluorescence-based enzymatic assays for galactose, which are associated with relatively high measurement uncertainty [46]. In contrast, uridine concentrations were quantified using HPLC [9] and therefore exhibited lower measurement noise. Accordingly, a lower measurement-to-process noise ratio is tuned for uridine compared to galactose. Because process noise is assumed constant over the culture duration, small fluctuations are introduced even during periods when uridine concentrations should remain at zero prior to feeding. This reflects a limitation of using fixed process noise and could be mitigated through adaptive tuning, although at the cost of increased filter complexity.

Across all feeding strategies, glucose accumulated to higher levels than predicted by the mechanistic model as seen in Figure 4 (m)-(p). This behaviour is consistent with the reduced cell growth observed experimentally when galactose and glucose are both supplied to the culture. When galactose is supplemented, carbon flux can be redirected toward central energy metabolism, reducing reliance on glucose as the primary carbon and energy source. Such shifts in substrate utilisation have been reported previously for this cell line [25] as well as other CHO cell systems [47]. The EnKF adjusted glucose trajectories accordingly, yielding substantially improved agreement with experimental measurements. Accurate glucose estimation is particularly important in this context, as glucose availability directly influences the synthesis rates of intracellular precursor pools that support glycosylation.

As shown in Figure 4 (q)-(t), the mechanistic model also consistently overpredicted extracellular glutamine concentrations across all conditions, likely reflecting limitations in parameter transferability to the new datasets. Measurement assimilation via the EnKF corrected these deviations and recovered trajectories that closely matched the observed profiles. This correction is especially relevant because the model assumes a direct relationship between extracellular and intracellular glutamine, with intracellular glutamine entering rate expressions governing the synthesis of NSD. Improved estimation of glutamine dynamics therefore directly enhances the reliability of inferred intracellular NSD concentrations and subsequent glycosylation predictions.

### 3.3 EnKF Estimation of Unmeasured Extracellular Asparagine

Asparagine plays a central metabolic role in mammalian cell culture, acting as a nitrogen donor for amino acid and nucleotide biosynthesis, supporting protein translation, and contributing to TCA cycle intermediates that sustain biomass growth and productivity [48]. Motivated by the strong empirical observability of asparagine identified in Figure 3(a), the EnKF is applied to infer its dynamics using only routinely available extracellular measurements. As shown in Figure 5, the filter reconstructs the asparagine trajectory with high accuracy, closely matching experimental measurements that are reserved exclusively for validation. In contrast, the open-loop mechanistic model fails to capture the observed dynamics, highlighting the limitations of fixed-parameter simulation under process variability.

**Figure 5:**
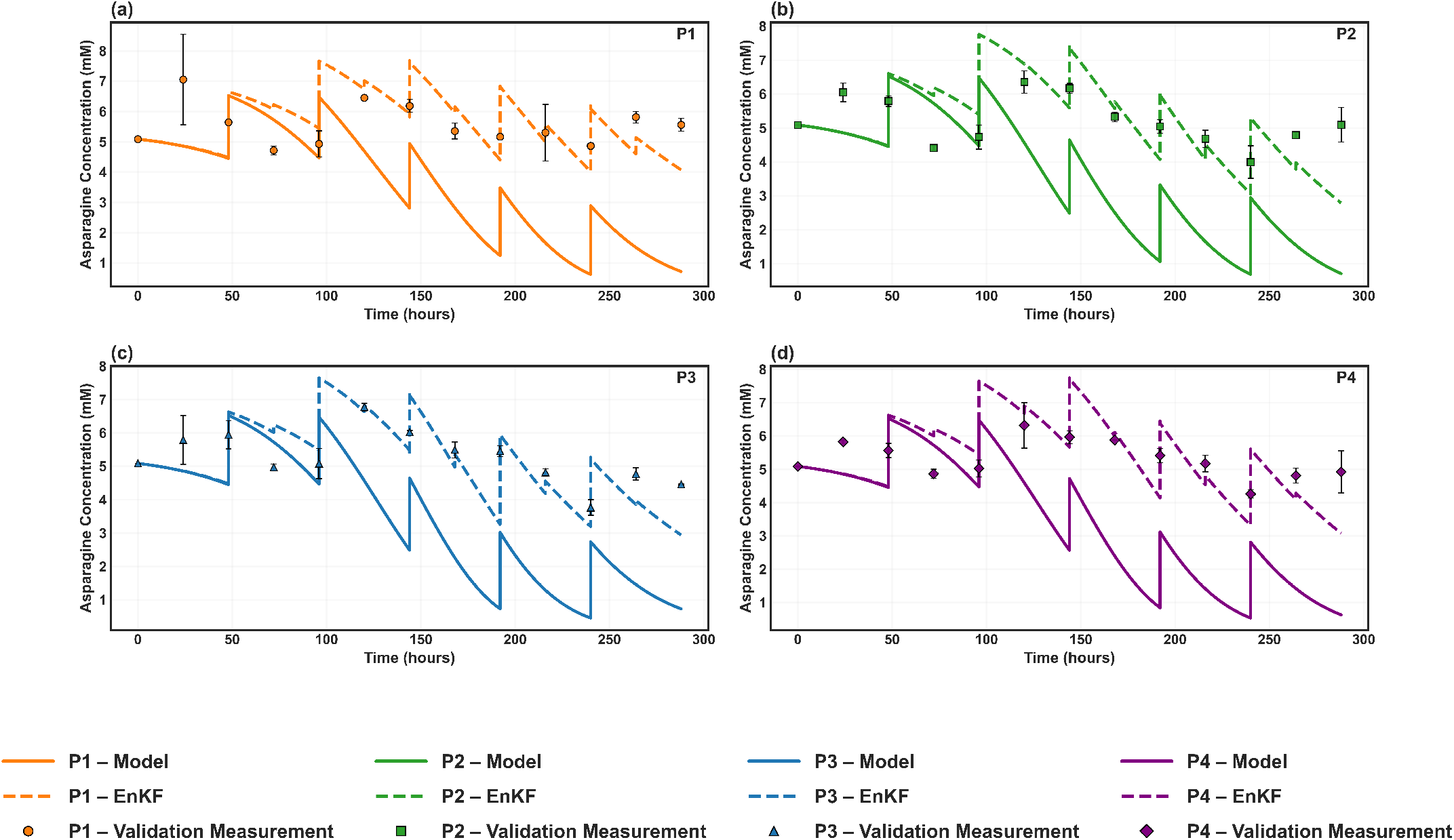
Comparison of asparagine trajectories predicted by the open-loop mechanistic model and estimated using the EnKF against experimental measurements. Only routinely available measurements are assimilated in the EnKF updates, including viable cell density, monoclonal antibody titre, galactose, uridine, glucose, glutamine, lactate, and ammonia. Asparagine measurements are used for validation only and are not assimilated by the EnKF.

### 3.4 EnKF Soft-Sensing of NSD

#### 3.4.1 Galactosylation precursor dynamics: UDP-Gal

UDP-Gal is the immediate intracellular precursor for terminal galactosylation of N-linked glycans [49]. It provides the mechanistic link between extracellular galactose and uridine supplementation and galactosylation, a key quality attribute of monoclonal antibodies [13, 50].

Across the four feeding strategies, the accuracy of UDP-Gal predictions is strongly influenced by how well extracellular galactose dynamics are captured. In the open-loop simulations, the mechanistic model systematically underpredicted galactose concentration as shown in Figure 4 (e)-(h), particularly in the later stages of culture. This mismatch propagated directly to the intracellular precursor pool, resulting in a pronounced underestimation of UDP-Gal toward the end of the process. Assimilation of galactose measurements through the EnKF substantially reduced this discrepancy. By correcting extracellular galactose trajectories, the filter enabled more reliable propagation of information into the intracellular precursor synthesis pathway, leading to improved reconstruction of UDP-Gal dynamics as shown in Figure 6 (a)-(d).

**Figure 6:**
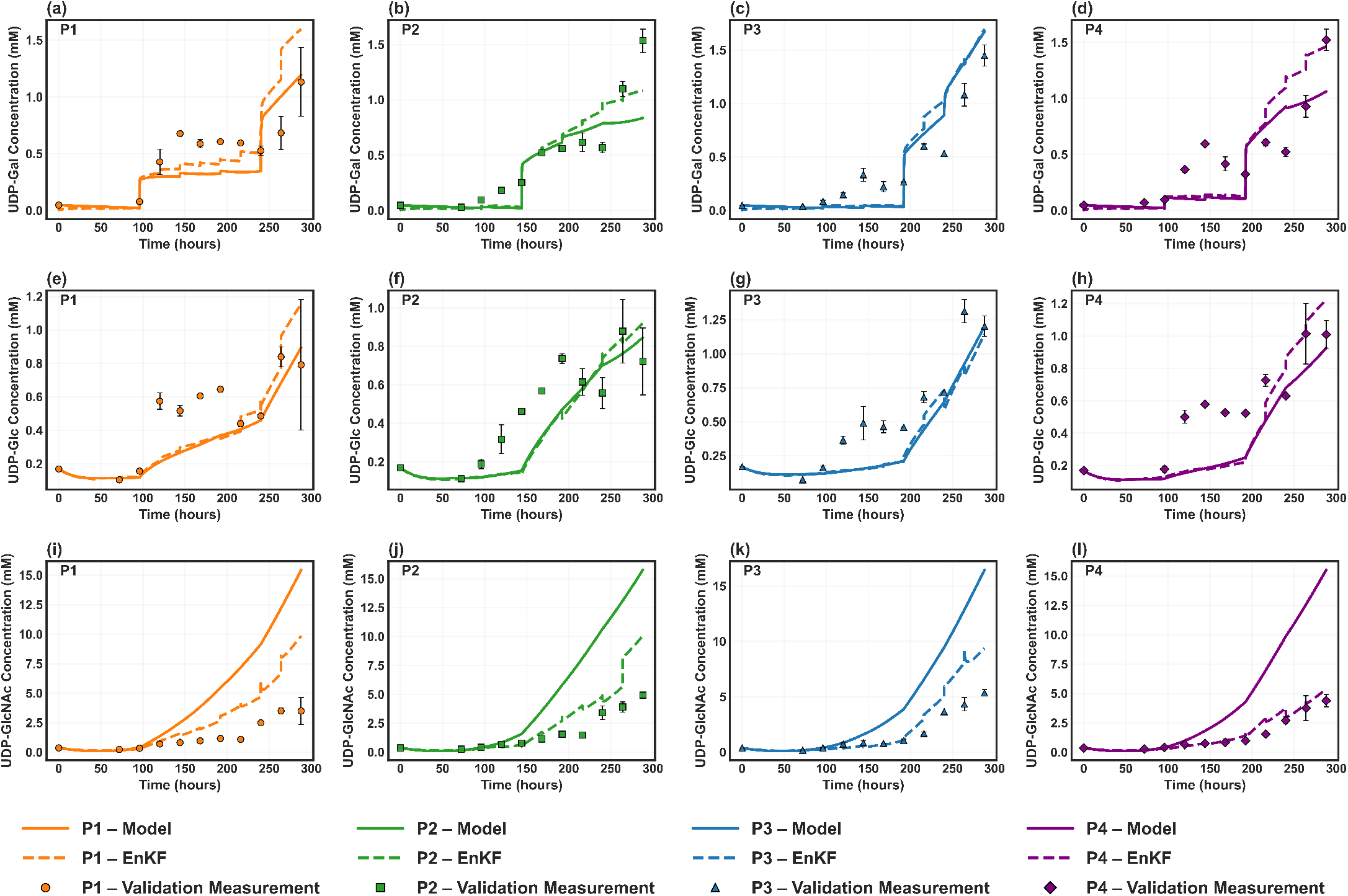
EnKF-based soft sensing of intracellular nucleotide sugar donors across four feeding strategies (columns). Rows correspond to UDP-Gal, UDP-Glc, and UDP-GlcNAc, respectively. Only routinely available measurements are assimilated in the EnKF updates, including viable cell density, monoclonal antibody titre, galactose, uridine, glucose, glutamine, lactate, and ammonia. NSD measurements are used for validation only, not used for EnKF updates.

#### 3.4.2 Central carbon-glycosylation coupling: UDP-Glc

UDP-Glc provides a direct link between central carbon metabolism and glycosylation and is primarily governed by extracellular glucose availability [51]. Under galactose supplementation, reduced glucose uptake leads to elevated extracellular glucose concentrations and altered intracellular precursor dynamics [47]. In this study, EnKF assimilation accurately reconstructed extracellular glucose profiles across all conditions as shown in Figure 4 (m)-(p), indicating that upstream carbon availability and extracellular variability are reliably captured. Despite this, UDP-Glc concentrations remained systematically underestimated during the mid-culture phase, indicating that the dominant source of mismatch lies within the intracellular NSD submodel rather than uncertainty in extracellular glucose dynamics.

Another factor to consider is the underlying reaction network, in which NSDs are tightly interlinked, particularly UDP-Glc and UDP-Gal, and which therefore dictates the model structure. This represents a structural limitation of the NSD module and underpins several of the observed parametric challenges.Parameterisation of UDP-Glc synthesis kinetics is challenged by sparse and uncertain calibration data, particularly during early culture when intracellular concentrations are low and large sample volumes are required to exceed analytical detection limits. These constraints reduce practical identifiability and allow calibration bias to persist. In addition, the current model assumes a direct correspondence between extracellular glucose and intracellular availability, neglecting transport and dynamic regulation of glycolytic flux associated with cellular energy status [52]. While the EnKF propagates corrected extracellular states into the intracellular network, it operates within a fixed NSD submodel and therefore cannot compensate for parameterisation or deficiencies in unmodelled dynamics of structurally unobservable NSD pathways.

#### 3.4.3 Branching precursor with strong inferability: UDP-GlcNAc

UDP-GlcNAc occupies a central position in *N*-linked glycosylation process, serving as the donor substrate for multiple branching and modification reactions [53, 54]. From a modelling and estimation perspective, UDP-GlcNAc is particularly important because it lies upstream of several intracellular species that are weakly coupled to extracellular measurements. In contrast to these downstream intermediates, UDP-GlcNAc synthesis is strongly linked to intracellular glutamine availability, which is informed by extracellular glutamine measurements that can be reliably assimilated using the EnKF. This structural positioning makes UDP-GlcNAc one of the most effective intracellular states for soft sensing, allowing measurement-driven corrections to propagate to less accessible regions of the network.

In open-loop simulations, the mechanistic model consistently overpredicted UDP-GlcNAc concentrations across all feeding strategies, with discrepancies increasing during mid-to late-culture phases. This behaviour is directly linked to overprediction of extracellular glutamine in the absence of measurement correction. Incorporation of EnKF updates substantially reduced this bias by assimilating glutamine measurements alongside other metabolites influencing growth, thereby refining both production and consumption fluxes associated with UDP-GlcNAc. The resulting trajectories closely matched experimental measurements across all four conditions.

The robustness of UDP-GlcNAc estimation is further supported by the quality of its underlying model fidelity. Compared with other NSDs, UDP-GlcNAc is chemically more stable and less prone to degradation during sample preparation, resulting in lower experimental variability [55]. These more reliable measurements informed the original parameterisation of the model, even though NSD data are not used during filtering. Consequently, the combination of strong extracellular coupling, accurate upstream correction, and higher model fidelity yields consistently reliable UDP-GlcNAc estimates in all validation scenarios.

## 4 Conclusions

The presented study demonstrates the application of EnKF as a soft-sensing framework for state estimation in mammalian cell culture processes. By integrating a mechanistic bioprocess model with routinely available extracellular measurements, the EnKF enables systematic treatment of uncertainty arising from measurement noise, model mismatch, and process variability. The framework was validated experimentally using independent fed-batch datasets operated under distinct galactose and uridine feeding strategies that span the edges of a constrained design space, providing the test of EnKF generalisation beyond the conditions used for model development.

For directly measurable extracellular states, the EnKF consistently corrected model bias while attenuating measurement noise, yielding estimated trajectories that better represent the underlying process dynamics than either open-loop model predictions or raw measurements alone. This demonstrates the value of ensemble-based data assimilation for improving state estimates in nonlinear bioprocess models, particularly under realistic operating conditions where neither models nor measurements are individually reliable.

A key focus of this work was the estimation of asparagine, an analytically challenging but biologically critical nutrient that strongly influences cell growth and is tightly coupled with other measurable extracellular states. Empirical observability analysis confirmed that asparagine is structurally inferable from available measurements, despite being unmeasured online. Consistent with this analysis, the EnKF accurately reconstructed asparagine trajectories across validation experiments, closely matching experimental measurements that were reserved exclusively for EnKF assessment. These results establish asparagine as a viable state for soft sensing in mammalian cell culture and illustrate how observability analysis can guide the selection of estimable states.

However, the estimation of NSD is fundamentally constrained by the structure of the biological system. NSDs are governed by a strictly feedforward information structure, whereby extracellular metabolites drive their synthesis, but NSD concentrations do not influence measurable outputs. This renders NSDs structurally unobservable, a limitation that cannot be overcome by estimator choice alone. Combined with the experimental challenges associated with reliable NSD measurements and intrinsic process variability, this has motivated alternative approaches that bypass explicit NSD estimation. For example, Luo *et al*. proposed a linearised discrete-time model that maps extracellular variables directly to glycan outcomes, enabling classical observability analysis and Kalman filtering under linear assumptions [56], while Zupke *et al*. developed empirical nonlinear correlations between culture conditions and glycan attributes to regulate high-mannose species in industrial systems [57]. Although effective in specific contexts, these approaches rely on empirical approximations and sacrifice mechanistic interpretability.

In this work, the EnKF retains a transparent, first-principles model structure while providing a numerically robust means of conditioning NSD dynamics on corrected extracellular states. Although the EnKF does not resolve structural unobservability, its ensemble-based uncertainty propagation enables plausible inference of NSD trajectories without imposing artificial linear sensitivity, making it more appropriate than Jacobian-based filters that require local observability for stable operation. The EnKF-derived NSD trajectories were validated against experimentally measured NSD concentrations that directly govern glycosylation, despite these measurements not being used in the filtering procedure.

Under the structural unobservability inherent to this biological system, the accuracy of NSD inference through EnKF depends on two key factors. First, strong coupling between extracellular metabolites and specific NSDs is essential for reliable propagation of information from measurable states. For instance, galactose with UDP-Gal, glucose with UDP-Glc, and glutamine with UDP-GlcNAc. NSDs that are downstream of other intracellular species and lack direct coupling to measured variables are correspondingly less reliable when none of the NSDs are observed. Second, model fidelity within the NSD subsystem plays a critical role. Unlike extracellular states, which can be continuously corrected through measurement updates, NSDs are inferred entirely through mechanistic coupling. In this study, UDP-GlcNAc was reconstructed most accurately, reflecting both strong coupling to extracellular glutamine and higher confidence in the parameterisation of its synthesis pathway. This highlights that soft sensing of intracellular precursors is viable when supported by both strong biological coupling and reliable mechanistic representation, even when model predictions deviate substantially under new operating conditions.

To summarise, this work addresses a long-standing challenge in upstream bioprocessing, where intracellular precursors influencing product quality are not available for real-time monitoring and decision-making. By inferring unmeasured intracellular states from mechanistic predictions constrained by extracellular data, the EnKF brings model-based insight closer to operational decision-making without increasing measurement complexity. Such soft-sensed variables can support earlier and more informed interventions, guide feed design and timing, and improve understanding of quality attribute sensitivity during process development. From a process systems perspective, EnKF-based soft sensing provides a transparent, uncertainty-aware framework aligned with Quality-by-Design principles, offering a practical pathway toward model-informed monitoring and, ultimately, adaptive control in biopharmaceutical manufacturing.

## 5 Data Availability

The datasets, code, and analysis scripts supporting the findings of this study are openly available in the GitHub repository at https://github.com/ly1815/EnKF-Soft-Sensing.

## 6 Acknowledgments

Three authors have no competing interest to declare. Author contributions are as follows. LY: Conceptualization, Methodology, Formal Analysis, Writing - Original Draft; ARC: Conceptualization, Supervision, Writing - Review and Editing; CK: Conceptualization, Supervision, Writing - Review and Editing. The authors acknowledge the use of ChatGPT 5.2 (OpenAI) for improving the readability and language of this work.

## A Mechanistic Model from Kotidis *et al*. 2019

### A.1 Metabolism Submodel

The Metabolism submodel provides a representation of extracellular cell growth, metabolism, and monoclonal antibody (mAb) synthesis. The formulation is structured into three coupled modules: (i) CHO cell growth and death kinetics,(ii) extracellular metabolite and amino acid dynamics, and (iii) antibody production. Although each module is defined separately, their interactions are governed through shared state variables and kinetic dependencies.

The model captures the influence of key carbon sources, nitrogen donors, metabolic by-products, and supplemented precursors on cell growth and productivity. Specifically, glucose, galactose, glutamine, lactate, ammonia, glutamate, asparagine, aspartate, and uridine are considered, based on their established relevance to CHO cell metabolism and product formation.

#### CHO cell growth and death

Cell culture volume dynamics are governed by the balance between feed addition and sampling withdrawal:

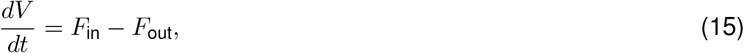

where *V* (L) denotes the culture volume, *t* (h) is time, *F*_in_ (L h^−1^) is the feed flow rate, and *F*_out_ (L h^−1^) is the sampling flow rate.

The viable cell density balance is expressed as:

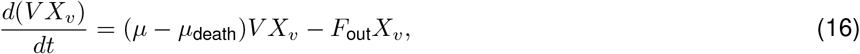

where *X*_*v*_ (cell L^−1^) is the viable cell density, *µ* (h^−1^) is the specific growth rate, and *µ*_death_ (h^−1^) is the specific cell death rate.

The specific growth rate is formulated as the product of a maximum growth capacity and regulatory terms capturing substrate limitation and metabolic inhibition:

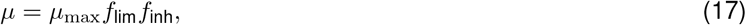

where *µ*_max_ (h^−1^) is the maximum specific growth rate.

Substrate limitation is attributed to extracellular glucose and asparagine concentrations:

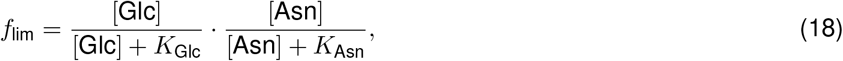

with [Glc] and [Asn] (mM) denoting extracellular glucose and asparagine concentrations, and *K*_Glc_ and *K*_Asn_ (mM) the corresponding Monod constants.

Growth inhibition is described through ammonia, lactate, and uridine:

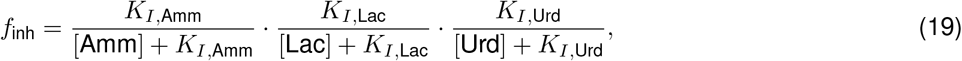

where [Amm], [Lac], and [Urd] (mM) are extracellular ammonia, lactate, and uridine concentrations, and *K*_*I*,Amm_, *K*_*I*,Lac_, and *K*_*I*,Urd_ (mM) are inhibition constants.

Cell death is assumed to be driven by ammonia and uridine accumulation:

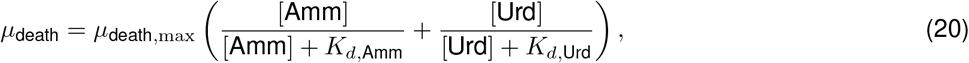

where *µ*_death,max_ (h^−1^) is the maximum death rate, and *K*_*d*,Amm_ and *K*_*d*,Urd_ (mM) are Monod constants for death induction.

#### CHO cell metabolism

Extracellular metabolite dynamics are described using a general material balance:

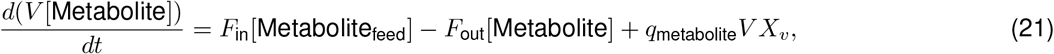

where [Metabolite] (mM) is the extracellular concentration, [Metabolite_feed_] (mM) is its concentration in the feed, and *q*_metabolite_ (mmol cell^*−1*^ h^−1^) is the specific production (positive) or consumption (negative) rate.

All biomass yield coefficients *Y*_*X/*metabolite_ are assumed constant throughout the culture to avoid overparametrisation.

#### Glucose

Glucose uptake supports glycolysis and pyruvate generation, but is modulated by the presence of galactose due to shared transport and phosphorylation pathways:

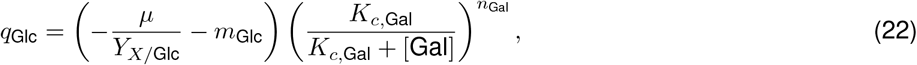

where *Y*_*X/*Glc_ (cell mmol^−1^) is the biomass yield on glucose, *m*_Glc_ (mmol cell^*−1*^ h^−1^) is the maintenance coefficient, *K*_*c*,Gal_ (mM) regulates galactose interference, and [Gal] (mM) is the extracellular galactose concentration.

The exponent *n*_Gal_ balances glucose and galactose contributions:

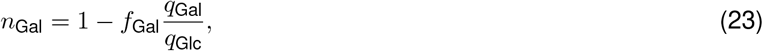

reflecting competitive uptake through shared transporters.

#### Glutamine

Glutamine dynamics include both growth-associated consumption and synthesis from ammonia:

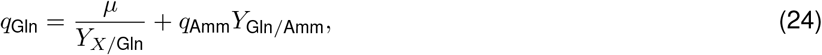

where *Y*_*X/*Gln_ (cell mmol^*−1*^) is the biomass yield on glutamine and *Y*_Gln*/*Amm_ (mmol mmol^*−1*^) represents glutamine synthesis via glutamine synthetase.

#### Lactate

Lactate production and consumption are described as:

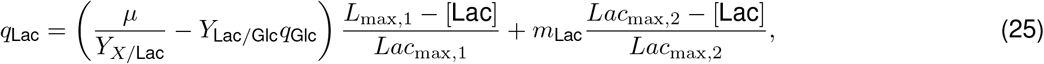

where *Y*_Lac*/*Glc_ (mmol mmol^*−1*^) is the lactate yield from glucose, *Lac*_max,1_ and *Lac*_max,2_ (mM) regulate lactate reutilisation, and *m*_Lac_ is the lactate maintenance coefficient.

#### Ammonia

Ammonia production arises from biomass formation and uridine metabolism:

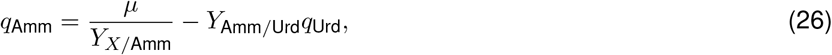

with *Y*_*X/*Amm_ (cell mmol^*−1*^) and *Y*_Amm*/*Urd_ (mmol mmol^*−1*^) denoting yield coefficients.

#### Glutamate

Glutamate uptake is assumed proportional to growth:

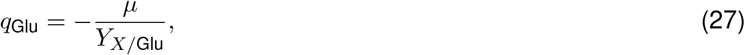

where *Y*_*X/*Glu_ (cell mmol^*−1*^) is the biomass yield on glutamate.

#### Galactose

Galactose consumption follows Monod kinetics:

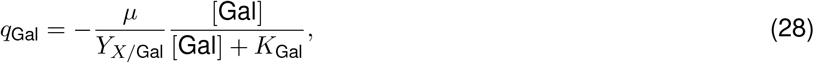

where *Y*_*X/*Gal_ (cell mmol^*−1*^) is the biomass yield on galactose and *K*_Gal_ (mM) is the Monod constant for galactose consumption.

#### Uridine

Similarly, uridine uptake is described by:

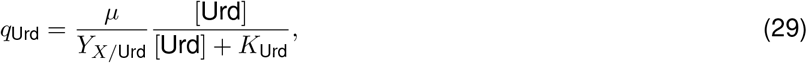

with *Y*_*X/*Urd_ (cell mmol^*−1*^) is yield of biomass on uridine and *K*_Urd_ (mM) is the Monod constant for uridine consumption.

#### Asparagine and Aspartate

Asparagine and aspartate are interconverted through enzymatic activity:

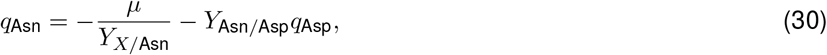

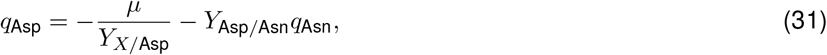

where *Y*_*X/*Asn_ and *Y*_*X/*Asp_ (cell mmol^*−1*^) are biomass yields, and *Y*_Asn*/*Asp_ and *Y*_Asp*/*Asn_ (mmol mmol^*−1*^) represent interconversion yields.

#### A.1.1 mAb synthesis

Monoclonal antibody production is described by

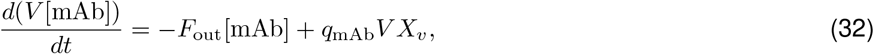

where [mAb] (mg*·*L^*−1*^) is the extracellular antibody concentration. The specific production rate is assumed to depend linearly on cell growth:

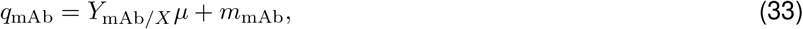

with *Y*_mAb*/X*_ denoting the growth-associated yield and *m*_mAb_ accounting for non-growth-associated production.

### A.2 NSD Submodel

The NSD submodel represents the intracellular metabolic network responsible for the synthesis and consumption of nucleotide sugar donors, which serve as activated co-substrates for *N-*linked glycosylation. The model accounts for seven intracellular NSD species, namely UDPGal, UDPGlc, UDPGalNAc, UDPGlcNAc, GDPMan, GDPFuc, and CMPNeu5Ac, and includes a total of sixteen enzymatic reactions.

Coupling between the NSD submodel and the Metabolism submodel is achieved through the specific cell growth rate, the specific monoclonal antibody production rate, and the extracellular concentrations of glucose, galactose, glutamine, and uridine. Reaction kinetics for NSD synthesis and interconversion are described using Monod-type saturation expressions. Based on experimental observations, intracellular nucleotide pools are assumed to be non-limiting and are therefore not explicitly included in the reaction rate expressions. To avoid excessive parameterization, intracellular concentrations of most metabolites and amino acids are not explicitly calculated. Glutamine constitutes an exception, as it directly participates in NSD synthesis reactions. Its intracellular concentration is approximated by a linear relationship with its extracellular concentration:

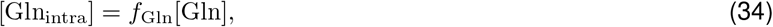

where [Gln_intra_] (mM) is the intracellular glutamine concentration and *f*_Gln_ is a dimensionless scaling factor.

#### A.2.1 NSD reaction network

A reduced NSD metabolic network is employed, capturing the dominant synthesis, conversion, and branching pathways relevant to CHO cell glycosylation. The formulation retains the mechanistic structure of the reported NSD biosynthetic pathways while limiting complexity to reactions that significantly influence cytosolic NSD availability. The reaction rates corresponding to the network are defined as follows:

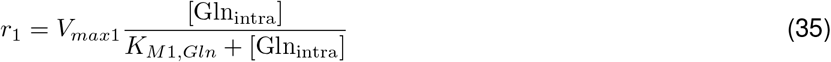

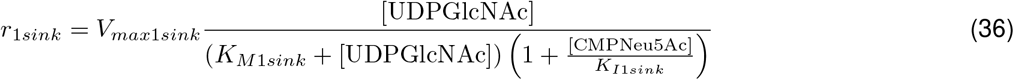

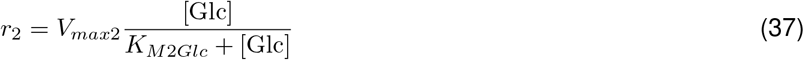

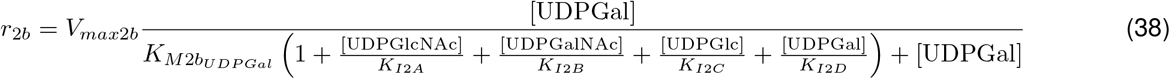

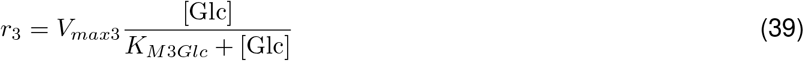

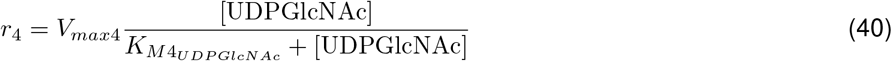

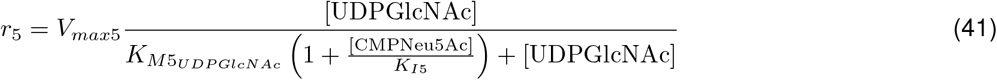

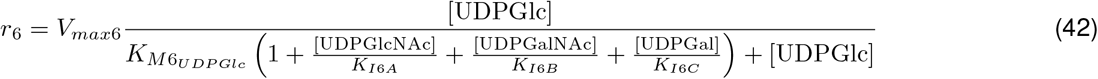

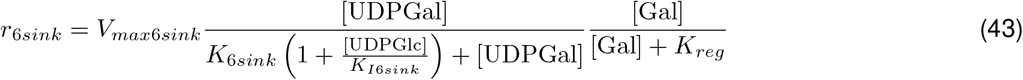

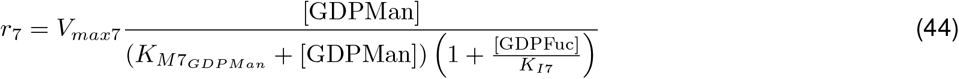

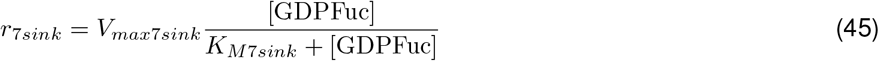

Here, 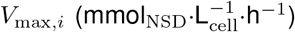 denotes the maximum rate of reaction *i, K*_*M*_ are saturation constants, and *K*_*I*_ represent inhibition constants. Sink reactions represent fluxes towards pathways not explicitly included in the reduced network but required to capture NSD depletion dynamics.

#### A.2.2 Effects of uridine and galactose supplementation

Additional reaction terms are introduced to account explicitly for the influence of uridine and galactose feeding on intracellular NSD levels. Uridine-dependent synthesis is described by

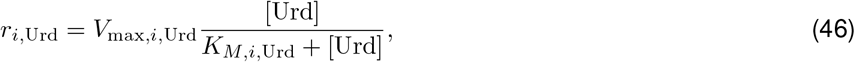

where *r*_*i*,Urd_ denotes the contribution of uridine to reaction *i*.

Galactose-enhanced UDPGal synthesis is represented as

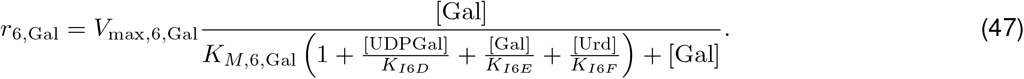

#### A.2.3 NSD material balances

The intracellular balance for each NSD species is given by

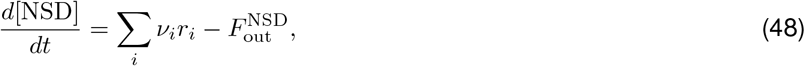

where *v*_*i*_ is the stoichiometric coefficient of reaction *i* for the considered NSD and 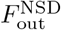 represents the net transport flux from the cytosol to the Golgi apparatus.

The transport flux is defined as

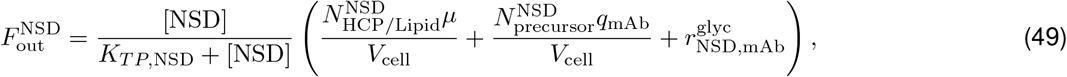

where *K*_*T P*,NSD_ is a transport saturation constant and *V*_cell_ is the cell volume.

The final term describes NSD consumption associated with monoclonal antibody glycosylation. To reduce computational cost while preserving mechanistic coupling with the Glycomodel, this term is approximated as

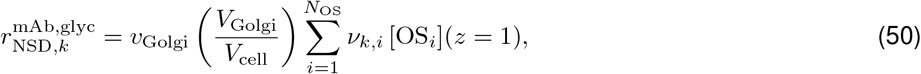

where *v*_Golgi_ (Golgi length·min^*−1*^) denotes the linear velocity of glycoprotein transport through the Golgi apparatus, *V*_Golgi_ (L) is the volume of the Golgi apparatus, and *V*_cell_ (L) represents the cellular volume. The coefficient 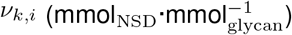 corresponds to the stoichiometric requirement of the *k*th NSD for the synthesis of the *i*th oligosaccharide, while [*OS*_*i*_](mmol_glycan_·L^*−1*^) denotes the concentration of the *i*th oligosaccharide at the exit of the Golgi apparatus (*z* = 1), as computed by the Glycomodel. The total number of oligosaccharide species included in the model is given by *N*_OS_, which is equal to 77 in the present study.

**Table S1:** Fitted parameters for the CHO cell growth, metabolism, and monoclonal antibody synthesis models.

| Parameter | Value | Unit |
| --- | --- | --- |
| $\mu_{max}$ | $6.50 \times 10^{-2}$ | $\text{h}^{-1}$ |
| $\mu_{death,max}$ | $1.50 \times 10^{-2}$ | $\text{h}^{-1}$ |
| $K_{Glc}$ | 14.04 | mM |
| $K_{Asn}$ | 2.62 | mM |
| $K_{I,Amm}$ | 3.17 | mM |
| $K_{I,Lac}$ | $1.00 \times 10^3$ | mM |
| $K_{I,Urd}$ | 41.09 | mM |
| $K_{d,Amm}$ | 14.28 | mM |
| $K_{d,Urd}$ | 27.86 | mM |
| $Y_{X/Glc}$ | $1.01 \times 10^9$ | $\text{cell} \cdot \text{mmol}^{-1}$ |
| $f_{Gal}$ | 0.35 | – |
| $m_{Glc}$ | $3.43 \times 10^{-11}$ | $\text{mmol} \cdot \text{cell}^{-1} \cdot \text{h}^{-1}$ |
| $K_{CGal}$ | 5.27 | mM |
| $Y_{X/Gln}$ | $4.64 \times 10^9$ | $\text{cell} \cdot \text{mmol}^{-1}$ |
| $Y_{Gln/Amm}$ | 0.10 | $\text{mmol} \cdot \text{mmol}^{-1}$ |
| $Y_{X/Lac}$ | $5.46 \times 10^7$ | $\text{cell} \cdot \text{mmol}^{-1}$ |
| $Y_{Lac/Glc}$ | 1.56 | $\text{mmol} \cdot \text{mmol}^{-1}$ |
| $Lac_{max1}$ | 21.20 | mM |
| $Lac_{max2}$ | 16.00 | mM |
| $m_{Lac}$ | $1.87 \times 10^{-10}$ | $\text{mmol} \cdot \text{cell}^{-1} \cdot \text{h}^{-1}$ |
| $Y_{X/Amm}$ | $2.36 \times 10^9$ | $\text{cell} \cdot \text{mmol}^{-1}$ |
| $Y_{Amm/Urd}$ | 2.00 | $\text{mmol} \cdot \text{mmol}^{-1}$ |
| $Y_{X/Glu}$ | $1.46 \times 10^{10}$ | $\text{cell} \cdot \text{mmol}^{-1}$ |
| $Y_{X/Gal}$ | $1.38 \times 10^8$ | $\text{cell} \cdot \text{mmol}^{-1}$ |
| $K_{Gal}$ | 18.23 | mM |
| $Y_{X/Urd}$ | $1.61 \times 10^9$ | $\text{cell} \cdot \text{mmol}^{-1}$ |
| $K_{Urd}$ | 7.00 | mM |
| $Y_{X/Asn}$ | $7.68 \times 10^8$ | $\text{cell} \cdot \text{mmol}^{-1}$ |
| $Y_{Asn/Asp}$ | 0.10 | $\text{mmol} \cdot \text{mmol}^{-1}$ |
| $Y_{X/Asp}$ | $3.59 \times 10^9$ | $\text{cell} \cdot \text{mmol}^{-1}$ |
| $Y_{Asp/Asn}$ | 0.13 | $\text{mmol} \cdot \text{mmol}^{-1}$ |
| $Y_{mAb,X}$ | 3.39 | $\text{pg} \cdot \text{cell}^{-1}$ |
| $m_{mAb}$ | $4.10 \times 10^{-1}$ | $\text{pg} \cdot \text{cell}^{-1} \cdot \text{h}^{-1}$ |

**Table S2:** Fitted parameters for the NSD submodel (Part I).

| Parameter | Value | Unit |
| --- | --- | --- |
| $f_{gln}$ | $2.00 \times 10^{-2}$ | N/A |
| $K_{M1,Gln}$ | 0.42 | mM |
| $K_{M1sink}$ | $4.00 \times 10^{-2}$ | mM |
| $K_{I1sink}$ | $1.21 \times 10^{-4}$ | mM |
| $K_{M2,Glc}$ | 78.12 | mM |
| $K_{M2,UDPGal}$ | $2.48 \times 10^{-2}$ | mM |
| $K_{I2A}$ | $1.05 \times 10^{-6}$ | mM |
| $K_{I2B}$ | 92.11 | mM |
| $K_{I2C}$ | $1.33 \times 10^{-2}$ | mM |
| $K_{I2D}$ | $2.66 \times 10^{-6}$ | mM |
| $K_{M3,Glc}$ | 50 | mM |
| $K_{M4,UDPGlcNAc}$ | 2.31 | mM |
| $K_{M5,UDPGlcNAc}$ | $2.70 \times 10^{-2}$ | mM |
| $K_{I5}$ | 1000 | mM |
| $K_{M6,UDPGlc}$ | $1.6 \times 10^{-2}$ | mM |
| $K_{I6A}$ | $1.10 \times 10^{-7}$ | mM |
| $K_{I6B}$ | 4.57 | mM |
| $K_{I6C}$ | $4.84 \times 10^{-6}$ | mM |
| $K_{6sink}$ | 0.13 | mM |
| $K_{I6sink}$ | 10 | mM |
| $K_{regulator}$ | $1.00 \times 10^{-5}$ | mM |
| $K_{M7,GDPMan}$ | 0.99 | mM |
| $K_{I7}$ | $1.64 \times 10^{-2}$ | mM |
| $K_{M7sink}$ | 8.88 | mM |
| $V_{max1,Urd}$ | 0.15 | $\text{mmol}_{\text{NSD}} \cdot \text{cell}^{-1} \cdot \text{h}^{-1}$ |
| $V_{max2,Urd}$ | $4.52 \times 10^{-2}$ | $\text{mmol}_{\text{NSD}} \cdot \text{cell}^{-1} \cdot \text{h}^{-1}$ |
| $V_{max4,Urd}$ | $1.27 \times 10^{-2}$ | $\text{mmol}_{\text{NSD}} \cdot \text{cell}^{-1} \cdot \text{h}^{-1}$ |
| $V_{max6,Urd}$ | 5.34 | $\text{mmol}_{\text{NSD}} \cdot \text{cell}^{-1} \cdot \text{h}^{-1}$ |
| $V_{max7sink}$ | 10.94 | $\text{mmol}_{\text{NSD}} \cdot \text{cell}^{-1} \cdot \text{h}^{-1}$ |

**Table S3:** Fitted parameters for the NSD submodel (Part II).

| Parameter | Value | Unit |
| --- | --- | --- |
| $K_{M1,Urd}$ | 6.08 | mM |
| $K_{M2,Urd}$ | 13.63 | mM |
| $K_{M4,Urd}$ | 6.25 | mM |
| $K_{M6,Urd}$ | 0.44 | mM |
| $V_{max1sink}$ | 25.49 | $\text{mmol}_{\text{NSD}} \cdot \text{cell}^{-1} \cdot \text{h}^{-1}$ |
| $V_{max6sink}$ | 7.30 | $\text{mmol}_{\text{NSD}} \cdot \text{cell}^{-1} \cdot \text{h}^{-1}$ |
| $V_{max6Gal}$ | 40.90 | $\text{mmol}_{\text{NSD}} \cdot \text{cell}^{-1} \cdot \text{h}^{-1}$ |
| $K_{M6Gal}$ | 0.60 | mM |
| $K_{I6D}$ | $1.00 \times 10^{-2}$ | mM |
| $K_{I6E}$ | 99.63 | mM |
| $K_{I6F}$ | $9.11 \times 10^{-4}$ | mM |
| $K_{TP,UDPGlc}$ | 0.90 | mM |
| $K_{TP,UDPGlcNAc}$ | 5.05 | mM |
| $K_{TP,UDPGal}$ | 7.15 | mM |
| $K_{TP,UDPGalNAc}$ | 11.06 | mM |
| $K_{TP,GDPMan}$ | 0.13 | mM |
| $K_{TP,GDPFuc}$ | 0.10 | mM |
| $K_{TP,CMPNeu5Ac}$ | 503.21 | mM |
| $V_{cell}$ | $1.12 \times 10^{-12}$ | L |
| $V_{Golgi}$ | $2.50 \times 10^{-14}$ | L |
| $MW_{mAb}$ | $165.17 \times 10^3$ | $\text{g}_{mAb} \cdot \text{mol}_{mAb}^{-1}$ |
| $N_{HCP/Lipids}^{UDPGlc}$ | $1.56 \times 10^{-12}$ | $\text{mmol}_{\text{NSD}} \cdot \text{cell}^{-1}$ |
| $N_{HCP/Lipids}^{UDPGlcNAc}$ | $1.25 \times 10^{-12}$ | $\text{mmol}_{\text{NSD}} \cdot \text{cell}^{-1}$ |
| $N_{HCP/Lipids}^{UDPGal}$ | $2.29 \times 10^{-12}$ | $\text{mmol}_{\text{NSD}} \cdot \text{cell}^{-1}$ |
| $N_{HCP/Lipids}^{UDPGalNAc}$ | $1.25 \times 10^{-12}$ | $\text{mmol}_{\text{NSD}} \cdot \text{cell}^{-1}$ |
| $N_{HCP/Lipids}^{GDPMan}$ | $3.54 \times 10^{-12}$ | $\text{mmol}_{\text{NSD}} \cdot \text{cell}^{-1}$ |
| $N_{HCP/Lipids}^{GDPFuc}$ | $1.40 \times 10^{-13}$ | $\text{mmol}_{\text{NSD}} \cdot \text{cell}^{-1}$ |
| $N_{HCP/Lipids}^{CMPNeu5Ac}$ | $1.85 \times 10^{-12}$ | $\text{mmol}_{\text{NSD}} \cdot \text{cell}^{-1}$ |
| $N_{precursor,UDPGlc}$ | $40.39 \times 10^{-6}$ | $\text{mmol}_{\text{NSD}} \cdot \text{mg}_{mAb}^{-1}$ |
| $N_{precursor,GDPMan}$ | $121.2 \times 10^{-6}$ | $\text{mmol}_{\text{NSD}} \cdot \text{mg}_{mAb}^{-1}$ |
| $N_{precursor,UDPGlcNAc}$ | $26.67 \times 10^{-6}$ | $\text{mmol}_{\text{NSD}} \cdot \text{mg}_{mAb}^{-1}$ |

## B Experiments

### B.1 Feeding Strategies of Datasets used for Model Calibration

**Table S4:** Galactose and uridine feed concentrations (mM) used in model calibration datasets.

| Experiment | Galactose in Feed C (mM) |  | Uridine in Feed C (mM) |  |
| --- | --- | --- | --- | --- |
|  | Day 4 | Day 8 | Day 4 | Day 8 |
| Control | 0 | 0 | 0 | 0 |
| 10G | 100 | 100 | 0 | 0 |
| 10G5U | 100 | 100 | 50 | 50 |
| 10G20U | 100 | 100 | 200 | 200 |
| 50G5U | 500 | 500 | 50 | 50 |

### B.2 Analytical Quantification

#### Quantification of viable cell density and antibody titre

Cell concentration and viability were measured using a Neubauer haemocytometer with trypan blue exclusion. Antibody titres in the culture supernatant were quantified using the BLItz® system with Dip and Read™ Protein A biosensors (Pall ForteBio, Portsmouth, UK).

#### Amino acid analysis and metabolites quantification

Extracellular concentrations of glucose, glutamine, glutamate, lactate, and ammonia were measured using the BioProfile analyser (NOVA Biomedical, MA). Extracellular galactose was measured using the Amplex™ Red Galactose/Galactose Oxidase Assay Kit (Life Technologies, Paisley, UK). Uridine concentration was quantified using the same method as the nucleotide sugar donor as described in the next section. Asparagine concentration was determined by HPLC Alliance system (Waters, Hertfordshire, UK) using the AccQ-Tag™ kit following the manufacturer’s protocol.

#### Nucleotide sugar donor analysis

A cell pellet containing 2 × 10^6^ cells was obtained by centrifugation at 100 g for 5 mins. The pellet was then extracted with 400 *µ*L of ice-cold 50% v/v acetonitrile (Sigma-Aldrich, Dorset, UK) and incubated on ice for 10 min. After centrifugation at 10,000 g for 5 min at 4 °C, the supernatant was collected and dried using a SpeedVac (Savant Inc., MI, USA). Dried extracts were resuspended in 150 *µ*L of distilled water, filtered through 0.22 *µ*m centrifuge tube filters (Sigma-Aldrich, Dorset, UK) for 5 min at 100 g, and stored at ^−^80°C until analysis to prevent NSD degradation. NSD concentrations were quantified by high-performance anion-exchange chromatography (HPAEC) on a Dionex CarboPac PA1 column (Dionex, USA) as described by del Val *et al*. [9], using 3 mM sodium hydroxide (E1) and 1.5 M sodium acetate in 3 mM sodium hydroxide (E2) as eluents.

## C Process and Measurement Covariance

Process covariances Σ^*x*^ were specified to define the relative uncertainty across model states and to initialise the ensemble spread. The ensemble-based state covariance, estimated from ensemble statistics, evolves dynamically at each prediction and update step. In contrast, the additive process noise covariance used during state propagation is prescribed and held constant, and is defined as *Q* = *k*_*Q*_Σ^*x*^. This formulation preserves the relative uncertainty structure across states while controlling the overall magnitude of injected process noise through a fixed scalar factor, *k*_*Q*_ = 2.0 *×* 10^−6^.

Measurement uncertainty is characterised by a diagonal covariance matrix *R*, defined only for the measured variables.

**Table S5:** Process and measurement covariance used for experimental validation of EnKF.

| | Process Covariance $\Sigma^x$ | Measurement Covariance $R$ |
| --- | --- | --- |
| Xv | $2.45 \times 10^{15}$ | $2.20 \times 10^{14}$ |
| mAb | $3.00 \times 10^2$ | $1.00 \times 10^1$ |
| Gal | 1.00 | $5.00 \times 10^{-3}$ |
| Urd | 1.00 | $1.00 \times 10^{-3}$ |
| Glc | $2.00 \times 10^1$ | $8.00 \times 10^{-2}$ |
| Amm | $1.63 \times 10^{-1}$ | $5.00 \times 10^{-3}$ |
| Gln | $4.00 \times 10^{-1}$ | $1.00 \times 10^{-4}$ |
| Lac | 1.16 | $4.00 \times 10^{-1}$ |
| Asn | $1.00 \times 10^{-2}$ | — |
| Glu | $4.80 \times 10^{-2}$ | — |
| UDP-Gal | $2.00 \times 10^{-1}$ | — |
| UDP-GalNAc | $1.00 \times 10^{-2}$ | — |
| UDP-Glc | $6.00 \times 10^{-2}$ | — |
| UDP-GlcNAc | $1.00 \times 10^{-2}$ | — |
| GDP-Man | $2.00 \times 10^{-4}$ | — |
| GDP-Fuc | $2.00 \times 10^{-4}$ | — |
| CMP-Neu5Ac | $2.00 \times 10^{-1}$ | — |

## D RMSE of EnKF and Model Prediction Compared with Experimental Data

**Table S6:** RMSE for extracellular metabolites and intracellular NSDs across datasets P1–P4.

| State (unit) | P1 |  | P2 |  | P3 |  | P4 |  |
| --- | --- | --- | --- | --- | --- | --- | --- | --- |
|  | Model | EnKF | Model | EnKF | Model | EnKF | Model | EnKF |
| Xv (cell $L^{-1}$) | $2.12 \times 10^9$ | $5.93 \times 10^8$ | $1.76 \times 10^9$ | $5.11 \times 10^8$ | $1.83 \times 10^9$ | $5.01 \times 10^8$ | $2.05 \times 10^9$ | $5.25 \times 10^8$ |
| mAb (mg $L^{-1}$) | 62.06 | 69.94 | 84.32 | 79.67 | 67.75 | 69.16 | 54.94 | 68.06 |
| Gal (mM) | 7.31 | 4.19 | 6.51 | 3.84 | 7.14 | 4.08 | 4.50 | 3.07 |
| Urd (mM) | 0.55 | 0.18 | 0.61 | 0.20 | 0.62 | 0.28 | 0.51 | 0.21 |
| Glc (mM) | 8.08 | 4.59 | 5.60 | 4.34 | 8.78 | 5.17 | 8.05 | 4.76 |
| Amm (mM) | 1.86 | 1.72 | 1.82 | 1.71 | 1.48 | 1.72 | 1.63 | 1.68 |
| Gln (mM) | 0.74 | 0.10 | 0.76 | 0.12 | 0.74 | 0.12 | 0.80 | 0.11 |
| Lac (mM) | 7.53 | 6.97 | 7.82 | 7.40 | 7.67 | 7.27 | 7.81 | 7.37 |
| Asn (mM) | 2.74 | 1.29 | 2.48 | 1.29 | 2.58 | 1.19 | 2.65 | 1.17 |
| UDP-Gal (mM) | 0.23 | 0.26 | 0.20 | 0.24 | 0.21 | 0.24 | 0.28 | 0.28 |
| UDP-Glc (mM) | 0.19 | 0.21 | 0.27 | 0.27 | 0.22 | 0.22 | 0.24 | 0.25 |
| UDP-GlcNAc (mM) | 5.47 | 2.48 | 7.64 | 4.06 | 4.90 | 1.75 | 5.21 | 0.62 |

